# Distinct bioenergetic features of human invariant natural killer T (iNKT) cells enable retained functions in nutrient-deprived states

**DOI:** 10.1101/2021.04.29.442021

**Authors:** Priya Khurana, Chakkapong Burudpakdee, Stephan A. Grupp, Ulf H. Beier, David M. Barrett, Hamid Bassiri

## Abstract

Invariant natural killer T (iNKT) cells comprise a unique subset of lymphocytes that are primed for activation and possess innate NK-like functional features. Currently, iNKT cell-based immunotherapies remain in early clinical stages, and little is known about the ability of these cells to survive and retain effector functions within the solid tumor microenvironment (TME) long-term. In conventional T cells (T_CONV_), cellular metabolism is linked to effector functions and their ability to adapt to the nutrient-poor TME. In contrast, the bioenergetic requirements of iNKT cells – particularly those of human iNKT cells – at baseline and upon stimulation are not well understood; neither is how these requirements affect cytokine production or anti-tumor effector functions. We find that unlike T_CONV_, human iNKT cells are not dependent upon glucose or glutamine for cytokine production and cytotoxicity upon stimulation with anti-CD3 and anti-CD28. Additionally, transcriptional profiling revealed that stimulated human iNKT cells are less glycolytic than T_CONV_ and display higher expression of fatty acid oxidation (FAO) and adenosine monophosphate-activated protein kinase (AMPK) pathway genes. Furthermore, stimulated iNKT cells displayed higher mitochondrial mass and membrane potential relative to T_CONV_. Real-time Seahorse metabolic flux analysis revealed that stimulated human iNKT cells utilize fatty acids as substrates for oxidation more than stimulated T_CONV_. Together, our data suggest that human iNKT cells possess different bioenergetic requirements from T_CONV_ and display a more memory-like metabolic program relative to effector T_CONV_. Importantly, iNKT cell-based immunotherapeutic strategies could co-opt such unique features of iNKT cells to improve their efficacy and longevity of anti-tumor responses.

## INTRODUCTION

Invariant natural killer T (iNKT) cells comprise a subset of innate-like T lymphocytes with TCR specificity for glycolipid antigens presented by the monomorphic, MHC I-like molecule CD1d (Bendelac et al., 2007). iNKT cells possess innate-like effector cell features, including rapid activation, cytokine secretion, and trafficking to tumor sites; as such, iNKT cells bridge innate and adaptive immune responses (Brennan et al., 2013; Matsuda et al., 2008). The presence of both circulating and intratumoral iNKT cells predicts more favorable tumor prognosis and survival in patients with several solid and liquid tumors (reviewed in Wolf et al., 2018), suggesting that these cells play central roles in cancer immunity. This notion is supported by a body of literature (reviewed in Altman et al., 2015) that demonstrate that iNKT cells engage in both direct anti-tumor cytotoxicity against CD1d-expressing tumors (Bassiri et al., 2014; Kawano et al., 1997) and modulate the activity of many other immune cells, including natural killer (NK) cells, CD8^+^ T cells (Carnaud et al., 1999; Crowe et al., 2002; Iyoda et al., 2018; Metelitsa et al., 2001; Mise et al., 2016; Smyth et al., 2002), and myeloid cells (Kitamura et al., 1999; Mussai et al., 2012; Song et al., 2009).

Recently, iNKT cells have begun to be utilized as a platform for cellular immunotherapy, as either adoptively-transferred cells (Exley et al., 2017; Kunii et al., 2009; Yamasaki et al., 2011) or chimeric antigen receptor (CAR)-transduced effectors directed against tumor antigens in lymphoma and solid tumor models (Heczey et al., 2014; Rotolo et al., 2018). While the studies published to date have demonstrated some efficacy, these trials are in early clinical stages and very little is understood about the basic cellular properties of iNKT cells that govern their ability to adapt to the tumor microenvironment (TME). Thus, there is a great need to better understand the metabolic properties of these cells in order to better inform the design of iNKT cell-based solid tumor immunotherapies in the future, particularly those that challenge existing conventional T cell (T_CONV_)-based therapies.

In T_CONV_, cellular metabolism is tightly linked to effector functions. Upon TCR stimulation, T_CONV_ undergo metabolic reprogramming as they differentiate from naïve to effector states, shifting from predominant use of oxidative phosphorylation (OXPHOS) to a preferential reliance on glycolysis and glutaminolysis to fuel substrate biogenesis and effector functions (Pearce et al., 2013). In the TME, the long-term functional capacity of T_CONV_ is minimized as they compete with tumor cells and tumor-supporting myeloid cells for limited glucose and glutamine (Chang et al., 2015; Ho et al., 2015). In contrast, memory T cells, which predominantly utilize fatty acid oxidation (FAO), are more persistent within the TME (Buck et al., 2016; Scharping et al., 2016; Sukumar et al., 2013). In addition, regulatory T cells (T_REG_) rely on OXPHOS and FAO (Michalek et al., 2011), allowing them to maintain immunosuppressive functions within the TME. Given that the metabolic profiles of T_CONV_ and other immune cells have been demonstrated to directly influence tumor progression, the use of therapies that modulate TME metabolism represent attractive treatment options for solid tumors.

In contrast to T_CONV_, however, little is known about iNKT cell metabolism and its link to key anti-tumor effector functions such as cytokine production and cytotoxicity. Unlike T_CONV_, iNKT cells do not have distinct differentiation states and exit the thymus primed for activation (D’Andrea et al., 2000); these functional differences may indicate a unique underlying metabolic phenotype. Indeed, murine iNKT cells have been demonstrated to depend predominantly on OXPHOS for survival (Kumar et al., 2019) and have also been shown to increase lipid biosynthesis upon activation, both *in vitro* and within the TME (Fu et al., 2020). Together, these data suggest that murine iNKT cells may have different bioenergetic profiles from T_CONV_, which could have significant consequences for their survival and function within the TME. While these studies have begun to elucidate the metabolic profiles of murine iNKT cells, a metabolic characterization of human peripheral blood iNKT cells and how it is linked to anti-tumor effector functions relative to T_CONV_ has not been determined previously.

In the present study, we sought to delineate the metabolic and functional properties of rested and stimulated human iNKT cells relative to T_CONV_ under both normal and nutrient-deplete conditions. Using peripheral blood-derived iNKT cells and matched T_CONV_ from healthy human donors, we demonstrate distinct bioenergetic requirements between iNKT cells and T_CONV_ for cytokine production and cytotoxicity after TCR stimulation. Specifically, we demonstrate that iNKT cells maintain effector functions in glucose- and glutamine-depleted conditions, and furthermore, utilize FAO metabolism to a greater extent than do T_CONV_. Our findings not only unveil novel bioenergetic features of primary human iNKT cells, but also suggest that iNKT cells may possess enhanced adaptability and longevity within the TME. Importantly, these features could be co-opted in the design of future iNKT cell-based solid tumor immunotherapies.

## MATERIALS AND METHODS

### Human Primary Immune Cell Purification

Healthy, de-identified human donor peripheral mononuclear blood cells (PBMC) and conventional T cells (T_CONV_) were purchased from the University of Pennsylvania Human Immunology Core under an institutional review board-approved protocol. To obtain sufficient yields of invariant natural killer (iNKT) cells, PBMC were plated in AIM V media (Gibco) containing 500ng/mL alpha-galactosylceramide (Cayman Chemicals; KRN7000) and 50U/mL recombinant IL-2 (PeproTech); on day 3-4 of culture, cells were fed with 10ng/mL recombinant IL-15 (BioLegend) and 10U/mL IL-2. On days 7-8 of expansion, iNKT cells were FACS-sorted from expanded PBMC (Vα24^+^ CD3^+^ cells) on BD Aria II or Aria Fusion instruments housed at the University of Pennsylvania and the Children’s Hospital of Philadelphia Flow Cytometry Core Facilities, respectively. In parallel, purified CD4^+^ and CD8^+^ T cells from matched donors were mixed at a 1:1 ratio to ensure equal composition of T_CONV_ populations.

### Cell Culture and Stimulation of Purified Lymphocytes

Purified iNKT and pooled T_CONV_ (1:1 CD4^+^:CD8^+^) populations were subject to either “rest” (low dose 30U/mL IL-2 only) or stimulation using Dynabeads Human T-Expander CD3/28 (ThermoFisher Scientific) at 1 million cells/mL at a ratio of 2 beads per cell for 48 hours. For all studies of rested and stimulated cells under normal conditions (transcriptional profiling and flow-based dyes), cells were cultured in AIM V media containing 10% FBS and 1% L-glutamine. For glucose deprivation studies, cells were rested and stimulated in complete RPMI 1640 media (Gibco) containing 10% dialyzed FBS and 1% L-glutamine supplemented with either 10mM glucose, 1mM glucose, or 0.1mM glucose (Corning). For glutamine deprivation studies, cells were plated in either complete RPMI media (containing 10% FBS and 1% L-glutamine at an approximate concentration of 4mM glutamine total), or glutamine-free RPMI (Gibco) containing 10% dialyzed FBS supplemented with 10mM glucose. For inhibition of glucose metabolism, 2-deoxy-D-glucose (Sigma cat # D6134) was added to cells at concentrations of 2mM or 20mM for 48 hours.

### Flow Cytometry

To sort iNKT cells, the following antibodies were used for staining: anti-Vα24-Jα18 (clone 6B11; BioLegend #342912), anti-CD3 (clone OKT3; BioLegend #317318). For staining of purified lymphocyte populations, cells were first stained with Zombie Aqua fixable live/dead exclusion dye per manufacturer’s instructions (BioLegend cat #423101), followed by surface staining in FACS buffer containing 2.5% FBS. For intracellular staining, cells were fixed and permeabilized using the Becton Dickinson Cytofix/Cytoperm kit, according to manufacturer’s instructions (BD Biosciences cat #554714) and stained with antibodies against granzyme B (clone QA16A02; BioLegend #372208) or mouse IgG1k isotype control (clone MOPC-21; BD Biosciences cat #556650), or Cpt1a (clone8F6AE9; Abcam cat# 171449) and rabbit IgG monoclonal isotype control (clone EPR25A; Abcam cat# 199091). Samples were run on a FACSVerse cytometer (BD Biosciences) and analyzed using FlowJo software (Tree Star Inc.).

### Mitochondrial Dye Staining

Purified rested and stimulated human iNKT cells and T_CONV_ were harvested at 48 hours for staining in either 200nM MitoTracker Green (Invitrogen cat #M7514) or 20nM tetramethylrhodamine, methyl ester (TMRM; Invitrogen cat #T668) in serum-free RPMI media for 45 minutes at 37°C per manufacturer’s protocol.

### RNA Purification and Quantitative Real-Time PCR

Total RNA was isolated from rested and stimulated iNKT cells and T_CONV_ using miRNeasy Mini kit per manufacturer’s protocol (Qiagen). RNA was either hybridized for NanoString transcriptional analysis or converted to cDNA for qPCR analysis. For gene expression analysis of *Ifng*, cDNA was synthesized from purified mRNA using High Capacity Reverse Transcriptase kit (Applied Biosystems) according to manufacturer’s protocol. Quantitative real-time PCR was performed on 7900HT Fast Real-Time PCR system (Applied Biosystems). Relative gene expression was calculated by normalizing delta Ct values for each target probe to *Actb* levels for each sample using the 2^-ΔCt^ method. The following TaqMan Gene Expression Assays (Life Technologies) were used: human *Ifng* (Hs_00989291_m1), human *Actb* (Hs01060665_g1).

### NanoString nCounter Gene Expression Profiling and Analysis

Transcriptional profiling of mRNA isolated from rested and stimulated human iNKT cells and T_CONV_ was performed using the nCounter SPRINT Profiler (NanoString Technologies). Briefly, per manufacturer’s instructions, 50ng of each RNA sample was hybridized for 18 hours at 65ºC with reporter and capture probe sets for the Human Metabolic Pathways panel (containing 768 genes across several annotated metabolic pathways and 20 internal reference genes). Hybridized RNA samples were then loaded onto nCounter SPRINT cartridge to run on SPRINT Profiler instrument. Gene expression analysis was conducted using NanoString nSolver 4.0 software. Genes with counts under 100 were eliminated from analysis. Heatmaps were generated using Morpheus (https://software.broadinstitute.org/morpheus).

### Cytokine Analysis

Supernatants from 48-hour rested and stimulated iNKT cells and T_CONV_ were assayed for cytokine levels of IFN-γ using human ELISA kit (Invitrogen cat #88-7316) following manufacturer’s protocol. Quantification of TNF-α and IL-4 cytokines in supernatants was performed using V-Plex Pro-Inflammatory Panel 1 Human Kit (Meso Scale Discovery, cat #K15049D). Assays were performed per manufacturer’s protocol and read and analyzed on a Meso Scale Discovery QuickPlex SQ120 instrument.

### Seahorse XF Metabolic Analysis

Real-time metabolic measurements of oxygen consumption rate (OCR) and extracellular acidification rate (ECAR) of iNKT cells and T_CONV_ from matched donors were obtained using XFe96 Extracellular Flux Analyzer (Seahorse Biosciences). On the day prior to assay, XF cartridge was hydrated, and XF tissue microplates were coated with Cell Tak (Corning cat #354240) per manufacturer’s protocol. On the day of the assay, 48-hour stimulated iNKT cells and T_CONV_ were washed and seeded at a density of 220,000 cells per well on pre-coated tissue microplates in XF RPMI assay media (pH 7.4) supplemented with 10mM glucose, 2mM L-glutamine, and 1mM pyruvate. Cells were spun down at 1500rpm for 3 minutes to facilitate adherence and placed in non-CO_2_ incubator for one hour prior to running assay. The Long-Chain Fatty Acid Substrate Oxidation kit (Agilent cat #103672-100) was utilized to probe differences in OCR upon injection with either vehicle (media only) or etomoxir (4μM) to inhibit long-chain fatty acid oxidation. Following three basal measurements of OCR and ECAR, cells were sequentially injected with 1.5μM oligomycin A (ATP synthase inhibitor; Agilent Technologies), 0.5μM FCCP (mitochondrial uncoupling agent; Agilent Technologies), and 0.5μM rotenone/antimycin A (mitochondrial complex I and III inhibitors; Agilent Technologies). After injection of oligomycin A, six readings were taken; after the following two sequential injections, three readings were taken. Maximal respiration was calculated as the difference between OCR upon FCCP injection and non-mitochondrial respiration (OCR upon rotenone and antimycin A injection). ATP production was calculated as the difference in OCR prior to and after oligomycin A injection.

## RESULTS

### Human iNKT cells maintain anti-tumor effector functions in glucose-depleted culture conditions relative to T_CONV_

To reliably obtain sufficient numbers of iNKT cells for our studies, we used populations of expanded, de-identified healthy human donor peripheral blood mononuclear cells (PBMC) and purified conventional T cells (T_CONV_; equal ratio of CD4^+^ and CD8^+^ cells) from matched donors (as per the schematic in **Supplemental Figure 1A**). To compare the metabolic and functional properties of human iNKT cells relative to T_CONV_ under identical conditions, each cell type was subjected to 48 hours of either rest (low-dose IL-2 only) or stimulation using anti-CD3/anti-CD28-coated microbeads.

We first investigated the dependency of human iNKT cells on glucose for anti-tumor cytokine production and cytotoxicity. Glucose is a limited nutrient within the TME and is rapidly metabolized by highly glycolytic tumor cells. Indeed, several prior studies have demonstrated that *in vitro* glucose depletion impairs the effector functions of T_CONV_ (Cham and Gajewski, 2005; Cham et al., 2008; Chang et al., 2013) and that reliance on glycolysis confers poorer persistence and survival within the TME (Bengsch et al., 2016; Scharping et al., 2016). A recent study suggested that mouse iNKT cells uptake less glucose than CD4^+^ T cells (Kumar et al., 2019), suggesting that they may be less reliant on glucose metabolism. To assess the requirement of glucose for human iNKT cell effector functions, iNKT cells and T_CONV_ were rested or stimulated in culture conditions containing either standard glucose (10mM) or depleted glucose (1mM or 0.1mM) concentrations for 48 hours. Intriguingly, iNKT cells were able to maintain levels of both *Ifng* mRNA and secreted IFN-γ upon stimulation in low glucose media (**Figure 1A-1D**). In contrast, T_CONV_ were sensitive to glucose depletion and demonstrated a dose-dependent decrease in *Ifng* transcripts and IFN-γ secreted protein levels. Strikingly, in 0.1mM glucose conditions, T_CONV_ displayed an 85% reduction in stimulation-induced *Ifng* mRNA and an over 70% reduction in IFN-γ protein secretion relative to 10mM glucose, while iNKT cells had no significant changes in IFN-γ levels (**Figure 1A-1D**). In addition to IFN-γ secretion, we also observed a similar trend in the secretion of additional cytokines, including TNF-α and IL-4 (**Supplemental Figure 2A-2B**), whereby iNKT cells did not rely on glucose for stimulation-induced secretion of these cytokines while T_CONV_ demonstrated dose-dependent decreases in TNF-α and IL-4 secretion with reduced glucose.

**Figure 1:**
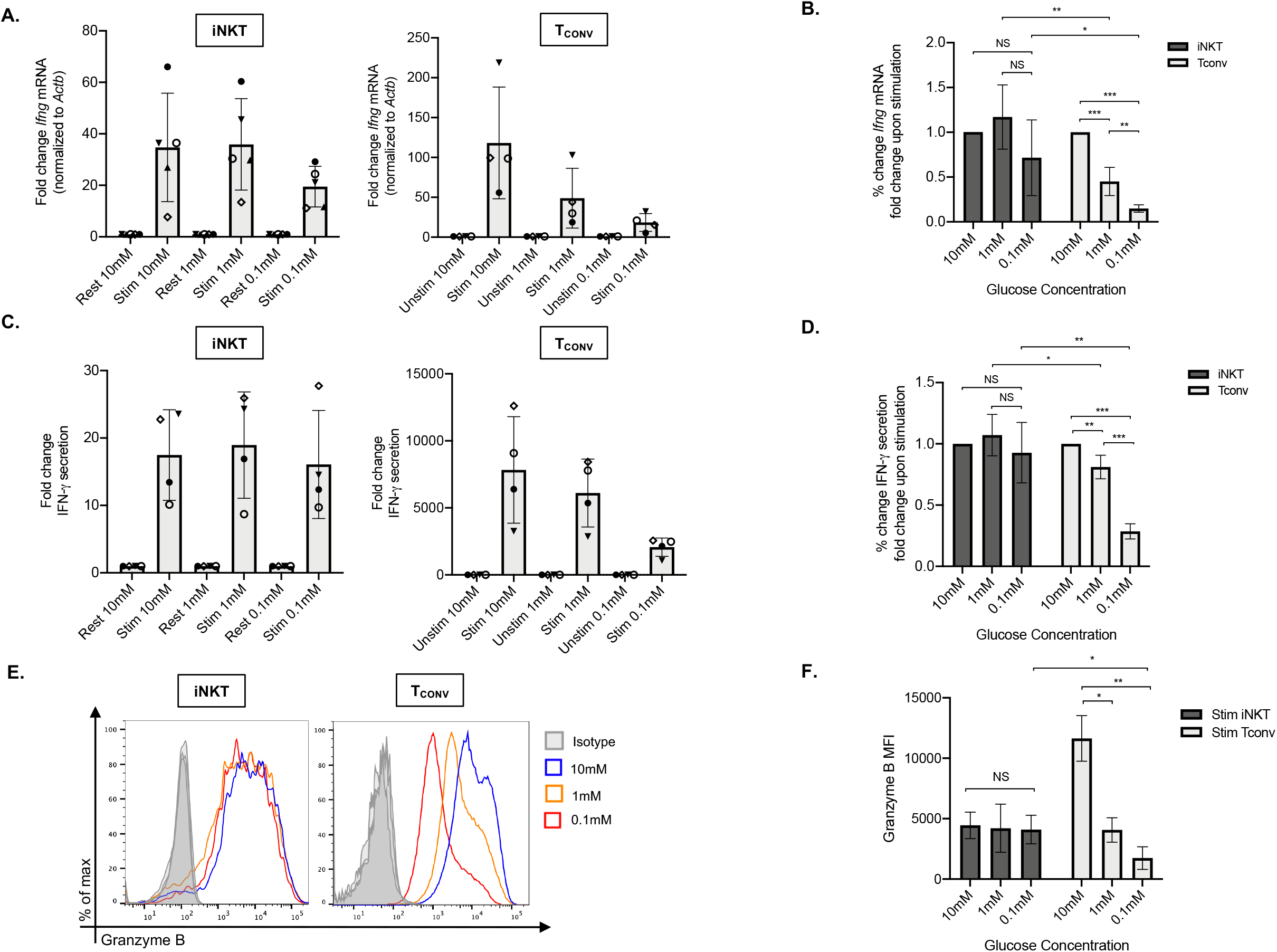
Human iNKT cells do not depend on glucose for anti-tumor effector functions relative to T_CONV_. Sorted PBMC-derived iNKT cells and T_CONV_ were rested or stimulated in RPMI media containing 10mM, 1mM, or 0.1mM glucose for 48 hours per schematic in Supplemental Figure 1. **(A)** mRNA expression of *Ifng* was determined for iNKT cells (left) and T_CONV_ (right) at 48 hours by qPCR (with values normalized to *Actb* expression). Fold change induction of *Ifng* upon stimulation relative to rest (iNKT) and unstimulated (T_CONV_) conditions displayed. Each symbol represents matched, independent human donor replicates. **(B)** Summary data of fold change in *Ifng* Ct upon stimulation (relative to 10mM glucose conditions) is depicted for iNKT and T_CONV_ from qPCR in (A). **(C)** Supernatants were collected from rested and stimulated iNKT cells (left) and T_CONV_(right) after 48 hours. IFN-γ levels were detected via ELISA and fold change upregulation upon stimulation relative to rest (iNKT) and unstimulated (T_CONV_) conditions displayed. Each symbol represents matched, independent human donor replicates. **(D)** Summary data of percent change in IFN-γ secretion fold change upon stimulation relative to 10mM glucose is depicted for iNKT and T_CONV_ from ELISA in (C). **(E)** Rested and stimulated iNKT cells and T_CONV_ from matched human donors were stained for intracellular Granzyme B or isotype control; histogram of live, stimulated iNKT cells and T_CONV_ representative of 4 matched, independent human donor samples. (F) Quantification of granzyme B mean fluorescence intensity (MFI) of stimulated iNKT cells and T_CONV_ normalized to isotype MFI indicated in bar graph. For all graphs, asterisks indicate statistical significance (*p<0.05, **p<0.01, ***p<0.001).

We next investigated the dependency of these cells on glucose for cytotoxicity by measuring levels of intracellular granzyme B, a surrogate for anti-tumor cytotoxic granule exocytosis. We found that stimulated iNKT cells maintain similar levels of intracellular granzyme B in glucose-deplete conditions, whereas T_CONV_ consistently displayed a significant reduction in granzyme B levels upon stimulation in lowered glucose concentrations (**Figure 1E**). Indeed, in 0.1mM glucose conditions, stimulated T_CONV_ granzyme B levels were reduced by approximately 85% relative to 10mM glucose, while stimulated iNKT cells displayed no significant difference in granzyme B with reduced glucose (**Figure 1F**). As an additional approach, we also treated iNKT cells and T_CONV_ with 2-deoxy-D-glucose (2-DG), a synthetic glucose analog that inhibits downstream glucose metabolism (Pajak et al., 2020). Although at a higher concentration of 2-DG (20mM), the effector functions of both iNKT cells and T_CONV_ were impaired, iNKT cells did still retain some level of IFN-γ production; however, at a lower concentration of 2-DG (2 mM), iNKT cells actually displayed moderately higher levels of cytokine production and cytotoxicity than in untreated conditions, while T_CONV_ were sensitive to glycolytic inhibition (**Supplemental Figure 3A-3C**). Collectively, these data further support the notion that iNKT cells are less glucose-dependent for effector functions than T_CONV_ and likely utilize alternate metabolic pathways upon stimulation.

### Human iNKT cells are less glycolytic than T_CONV_

The striking differences in the sensitivity of iNKT cells and T_CONV_ to glucose depletion suggest a potential underlying difference in glycolytic metabolism. While T_CONV_ upregulate glycolysis upon stimulation, the metabolic activity of human iNKT cells is unknown. Using a NanoString probe set with over 700 curated transcripts for genes involved in cancer immunology and metabolism, we assayed changes in the mRNA expression of metabolic genes in rested and stimulated PBMC-derived iNKT cells and matched T_CONV_ from 8 independent donors. In contrast to T_CONV_, which upregulated glycolytic pathway enzyme transcripts upon stimulation, iNKT cells only upregulated a small subset of the glycolytic genes upon stimulation, and to a lesser extent than T_CONV_ (**Figure 2A**). Indeed, of the 14 glycolytic genes probed, 9 were significantly differentially expressed between stimulated T_CONV_ and stimulated iNKT cells, and of these 9 genes, 7 were significantly higher in T_CONV_: *Hk2, Ldha, Ldhb, Aldoa, Eno1, Gapdh*, and *Pdha1* (**Figure 2B and Table 1**). Each of these genes encode key enzymes throughout the glycolysis pathway that also fuel additional biosynthetic pathways.

**Figure 2:**
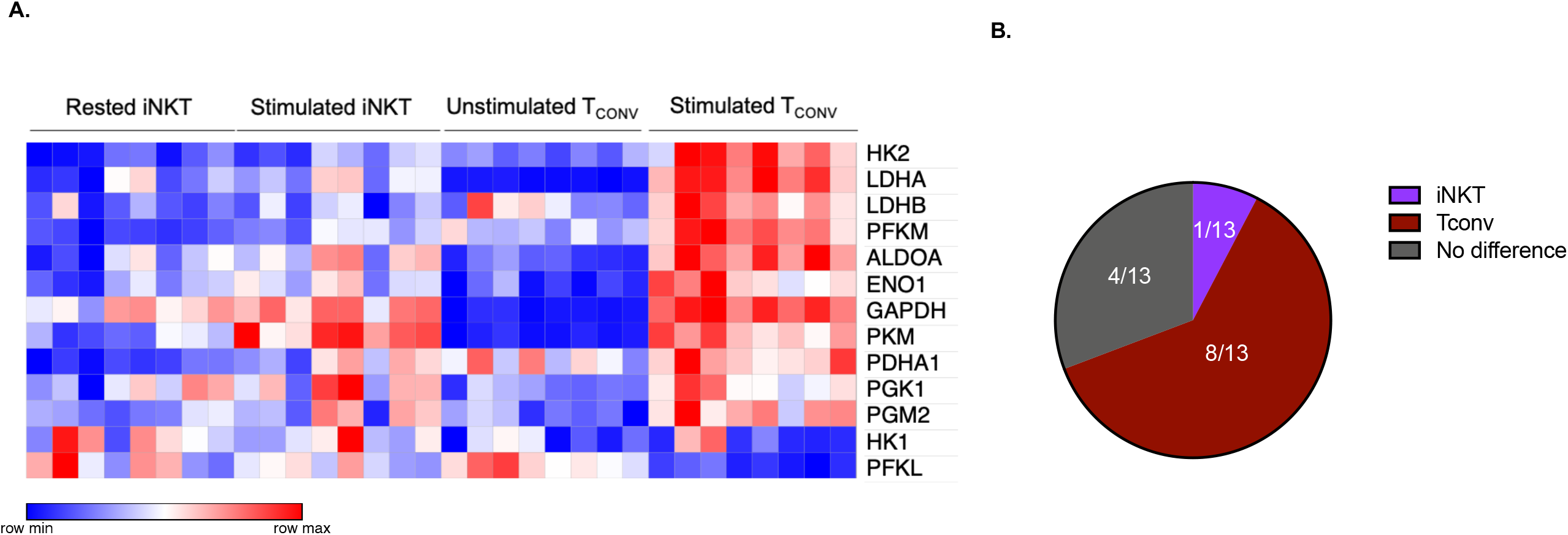
Human iNKT cells are less glycolytic than T_CONV_. **(A)** Heatmap of 8 independent, healthy human donor rested and stimulated iNKT cells and matched T_CONV_ (processed per schematic in Supplemental Figure 1) transcriptional profiles for genes in the glycolysis pathway included in the NanoString nCounter Human Metabolic Pathways probe set. Genes with counts under 100 were eliminated from analysis. Coloring indicates relative expression of each gene, from low (blue) to high (red). Heatmap generated on Morpheus. **(B)** Pie chart graphically displaying proportion of glycolysis genes significantly higher in each stimulated cell subset.

**Table 1:** Glycolysis pathway gene set expression in stimulated iNKT cells and T_CONV_.

| Gene name | p-value | q-value | Higher expressing cell subset |
| --- | --- | --- | --- |
| PFKL | 1.8003E-05 | 1.6498E-04 | iNKT |
| PFKM | 2.7224E-05 | 1.6498E-04 | T<sub>CONV</sub> |
| HK2 | 7.0098E-05 | 2.8319E-04 | T<sub>CONV</sub> |
| LDHA | 1.2875E-04 | 3.9011E-04 | T<sub>CONV</sub> |
| LDHB | 2.0950E-04 | 4.2319E-04 | T<sub>CONV</sub> |
| ALDOA | 6.6621E-03 | 1.0093E-02 | T<sub>CONV</sub> |
| ENO1 | 1.8666E-02 | 2.1844E-02 | T<sub>CONV</sub> |
| GAPDH | 1.9826E-02 | 2.1844E-02 | T<sub>CONV</sub> |
| PDHA1 | 2.7659E-02 | 2.7936E-02 | T<sub>CONV</sub> |
| PGM2 | 7.5981E-02 | 6.5778E-02 | T<sub>CONV</sub> |
| HK1 | 1.2941E-01 | 1.0457E-01 | iNKT |
| PKM | 2.0838E-01 | 1.5785E-01 | iNKT |
| PGK1 | 9.5339E-01 | 6.0816E-01 | iNKT |

In T_CONV_, the transcription factor Myc is a master regulator required for initiation and maintenance of glycolytic metabolic reprogramming after TCR stimulation (Wang et al., 2011). In iNKT cells, the role of Myc, particularly upon activation, has not been previously examined. Consistent with reduced glycolytic reprogramming in stimulated iNKT cells, we also found that iNKT cells have significantly lower upregulation of Myc pathway genes than stimulated T_CONV_ (**Supplemental Figure 4A-4B and Supplemental Table 1**). This difference in transcription of genes downstream of Myc signaling could suggest a mechanistic difference in the metabolic regulation of these two cell types.

Together, both the glycolysis and Myc pathway gene expression data suggest that human iNKT cells, in comparison to T_CONV_, employ distinct metabolic pathways from T_CONV_ upon TCR stimulation. Importantly, iNKT cells’ lack of dependence on glycolysis could be advantageous in the context of the TME, whereby iNKT cells may be able to maintain superior anti-tumor effector functions in glucose-diminished conditions in which T_CONV_ are at a disadvantage.

### Human iNKT cells are less sensitive to glutamine depletion than T_CONV_ for maintaining effector functions

Given that stimulated human iNKT cells were not dependent on glucose metabolism for effector functions, we wondered if they instead utilize glutamine as an alternative metabolic substrate to fuel cytokine production and cytotoxicity. Through glutaminolysis, glutamine is metabolized into α-ketoglutarate, which directly enters the TCA cycle to eventually yield ATP via oxidative phosphorylation (OXPHOS). Upon TCR activation, T_CONV_ increase both glutamine and glucose uptake and metabolism in order to fuel effector functions (Carr et al., 2010; Nakaya et al., 2014; Wang et al., 2011). In contrast, the role of glutamine has not yet been elucidated in iNKT cells. To investigate whether glutamine is required for iNKT cell effector functions, we rested and stimulated matched human iNKT and T_CONV_ in either complete or glutamine-free media using the schema described in Supplementary Figure 1. Upon stimulation in glutamine-deplete conditions, iNKT cells display a moderate decrease in *Ifng* mRNA expression of ∼40% of those stimulated in complete media, while T_CONV_ reduced stimulation-induced *Ifng* mRNA expression by ∼60% in glutamine-free conditions relative to complete media (**Figure 3A-3B**). Strikingly, IFN-γ secretion was not altered in iNKT cells stimulated in the absence of glutamine, whereas it was reduced by ∼90% in T_CONV_ (**Figure 3C-3D**). These data imply a differential reliance on glutamine for cytokine production between these cell types. We also observed preserved secretion of additional cytokines (TNF-α and IL-4) by iNKT cells stimulated in glutamine-deplete conditions (**Supplemental Figure 5A-5B**). Furthermore, while glutamine depletion almost entirely abrogated T_CONV_ intracellular granzyme B levels, iNKT cells were able to retain some granzyme B production even when stimulated in the absence of glutamine – measuring ∼65% of that of levels in normal conditions (**Figure 3E-3F**). Thus, this indicates that unlike T_CONV_, glutamine is not required for iNKT cell cytotoxicity. Collectively, our data demonstrate that iNKT cells are able to maintain their effector functions in both glutamine-deplete and glucose-deplete media conditions, which may have important consequences for their ability to exert anti-tumor immunity within the nutrient-deplete TME.

**Figure 3:**
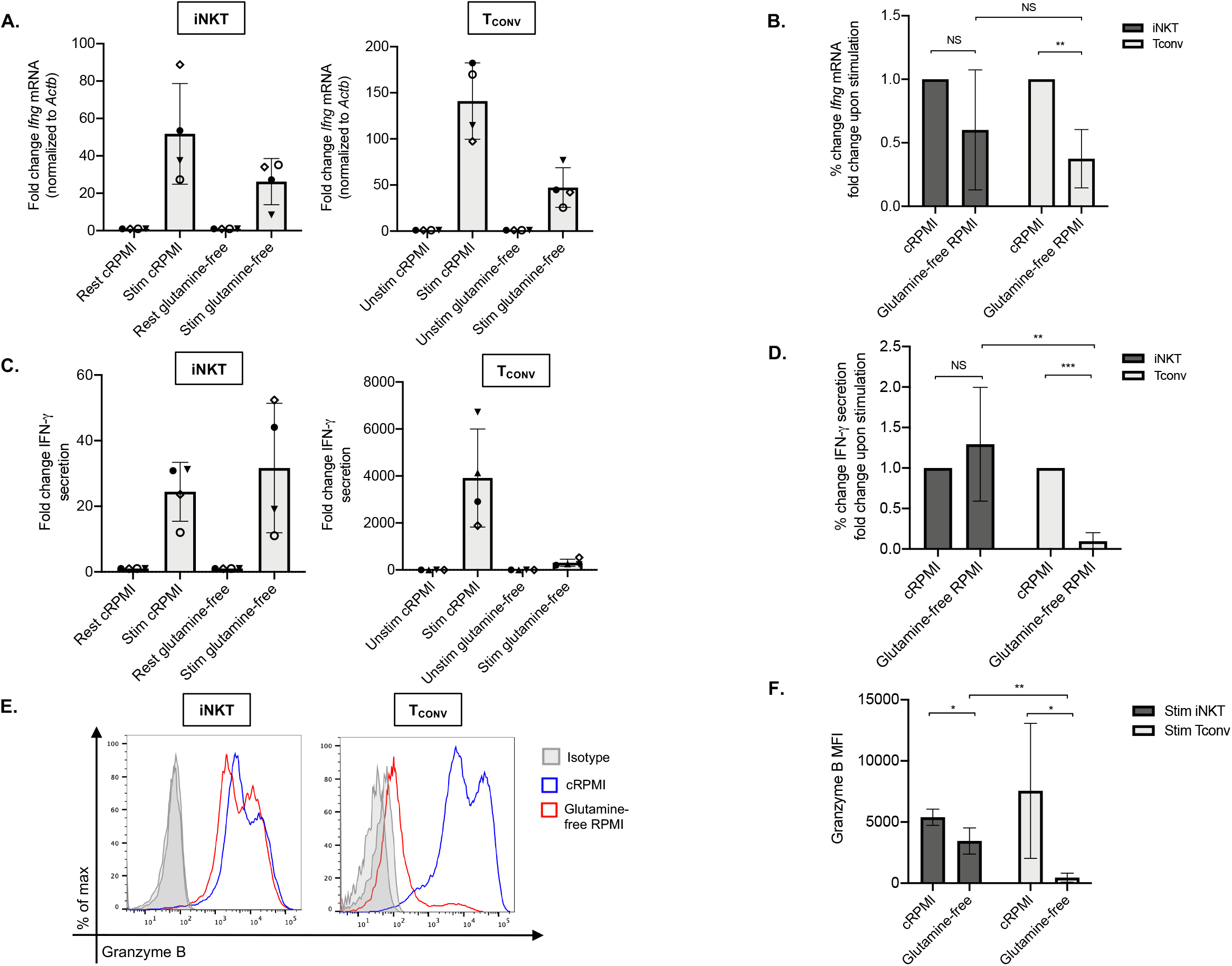
Human iNKT cells are less reliant on glutamine for anti-tumor effector functions than T_CONV_. Sorted PBMC-derived iNKT cells and T_CONV_ were rested or stimulated in either complete RPMI (10% FBS, 1% L-glutamine) or glutamine-free RPMI media for 48 hours per schematic in Supplemental Figure 1. **(A)** mRNA expression of *Ifng* was determined for iNKT cells (left) and T_CONV_ (right) at 48 hours by qPCR (with values normalized to *Actb* expression). Fold change induction of *Ifng* upon stimulation relative to rest (iNKT) and unstimulated (T_CONV_) conditions displayed. Each symbol represents matched, independent human donor replicates. **(B)** Summary data of fold change in *Ifng* Ct upon stimulation (relative to cRPMI condition) is depicted for iNKT and T_CONV_ from qPCR in (A). **(C)** Supernatants were collected from rested and stimulated iNKT cells (left) and T_CONV_ (right) after 48 hours. IFN-γ levels were detected via ELISA and fold change upregulation upon stimulation relative to rest (iNKT) and unstimulated (T_CONV_) conditions displayed. Each symbol represents matched, independent human donor replicates. **(D)** Summary data of percent change in IFN-γ secretion fold change upon stimulation relative to cRPMI is depicted for iNKT and T_CONV_ from ELISA in (C). **(E)** Rested and stimulated iNKT cells and T_CONV_ from matched human donors were stained for intracellular Granzyme B or isotype control; histogram of live, stimulated iNKT cells and T_CONV_ representative of 4 matched, independent human donor samples. **(F)** Quantification of granzyme B mean fluorescence intensity (MFI) of stimulated iNKT cells and T_CONV_ normalized to isotype MFI indicated in bar graph. For all graphs, asterisks indicate statistical significance (*p<0.05, **p<0.01, ***p<0.001).

### Human iNKT cells have a “memory-like” metabolic phenotype

Memory T cells that develop after initial antigenic activation are primed for rapid reactivation upon secondary antigenic encounter. To allow for greater self-renewal capacity and longevity, memory T cells are metabolically adapted to possess altered mitochondrial morphology with enhanced spare respiratory capacity and a predominant reliance on oxidative and lipid metabolism (van der Windt et al., 2012, 2013). We postulated that human iNKT cells may be “memory-like” in their metabolism. iNKT cells are poised for activation and express memory-like phenotypic markers, including CD62L, CCR7, and CD45RO (Baev et al., 2004; D’Andrea et al., 2000). Furthermore, murine NKT cells have been shown to depend on OXPHOS for survival, proliferation, and effector functions relative to CD4^+^ T cells (Kumar et al., 2019). Collectively, it also appears that iNKT cells do not have a distinct differentiation hierarchy of naïve, effector, and memory states, further suggesting that their underlying metabolic program may be unique from T_CONV_.

We first investigated mitochondrial parameters by flow cytometric dyes. Specifically, we utilized MitoTracker Green, which provides a measure of mitochondrial mass, as well as tetramethylrhodamine methyl ester perchlorate (TMRM), a cell permeable dye that accumulates in active mitochondria and serves as an indicator of mitochondrial membrane potential. We found that both resting and stimulated human iNKT cells have significantly higher mitochondrial mass (**Figure 4A**) and mitochondrial membrane potential (**Figure 4B**) relative to unstimulated and stimulated T_CONV_. Together, this may imply greater mitochondrial activity within iNKT cells relative to T_CONV_. Furthermore, NanoString transcriptional profiling of these cells revealed that resting and stimulated iNKT cells displayed significantly higher expression of several fatty acid oxidation (FAO) enzyme transcripts than stimulated T_CONV_ (**Figure 4C and Supplementary Table 2**). These genes include *Acaa2*, a mitochondrial enzyme involved in beta-oxidation of fatty acids into acetyl CoA, *Acat1* and *Acat2*, which convert ketones into acetyl-CoA, and *Acox1*, which also catalyzes beta-oxidation of fatty acids. One of the most striking differences was in the expression of *Cpt1a*, which encodes the rate-limiting enzyme of FAO that transports long-chain fatty acids into the mitochondria to be metabolized. Notably, *Cpt1a* was among the top 25 genes significantly higher in stimulated iNKT cells than stimulated T_CONV_, underscoring the potential importance of this enzyme for human iNKT cells. Indeed, in support of this transcriptional data, we also find that stimulated iNKT cells possess significantly higher levels of intracellular Cpt1a protein than rested iNKT cells and stimulated T_CONV_, as assessed by intracellular flow cytometry (**Figure 4D**). These data suggest that iNKT cells may predominantly utilize FAO metabolism upon stimulation, which could represent a key metabolic difference from T_CONV_.

**Figure 4:**
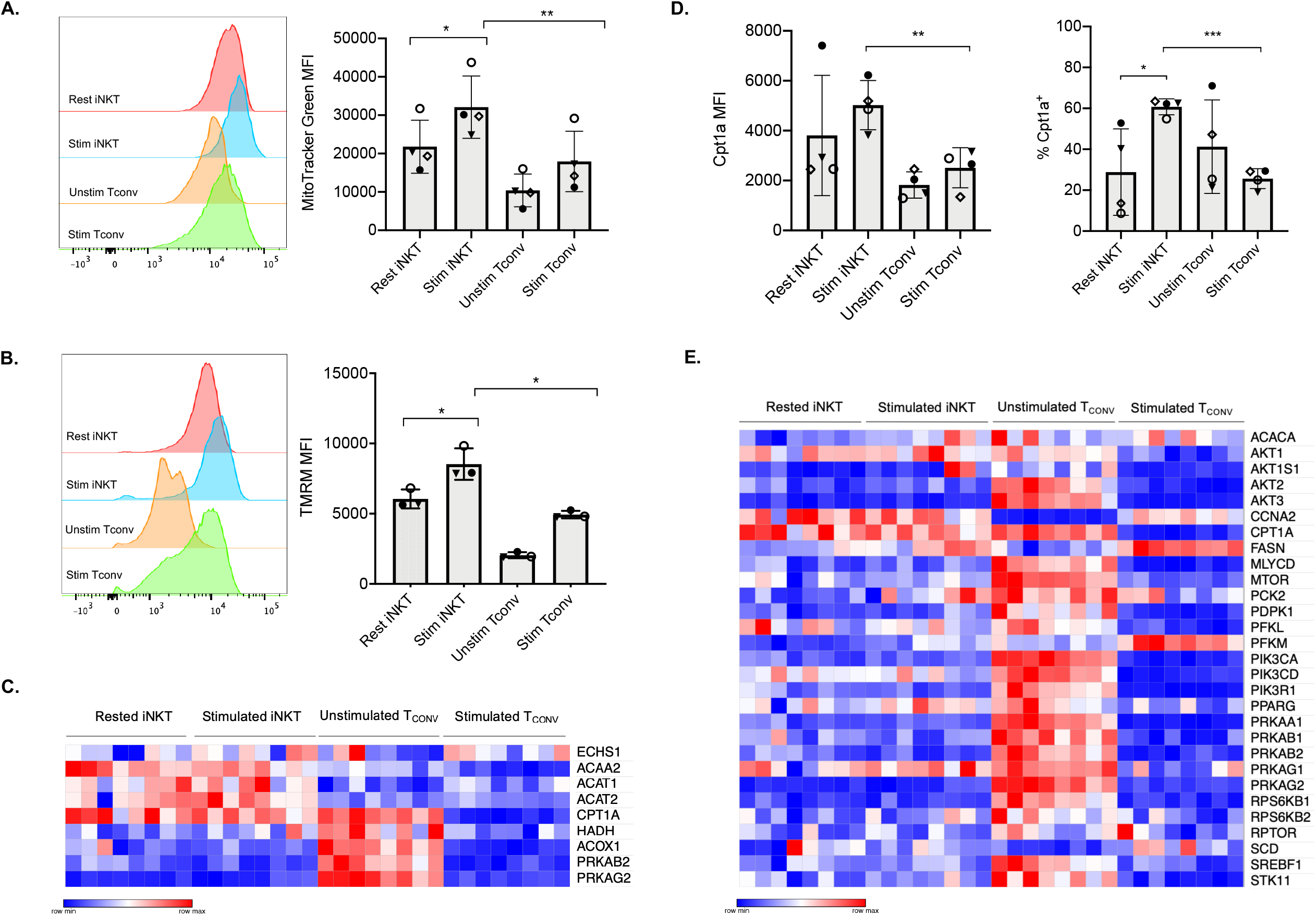
Human iNKT cells have a “memory-like” metabolic phenotype comprised of enhanced mitochondrial metabolism and FAO. **(A)** Histograms (left) displaying mean fluorescence intensity (MFI) of MitoTracker Green emission (FITC channel) in purified rested and stimulated iNKT cells and T_CONV_ from matching human donor samples. (Right) Bar graphs of MFIs. Symbols represent independent, matched human donors. **(B)** Histograms (left) displaying mean fluorescence intensity (MFI) of TMRM emission (PE channel) in purified rested and stimulated iNKT cells and T_CONV_ from matching human donor samples. (Right) Bar graphs of MFIs. Symbols represent independent, matched human donors. **(C)** Heatmap of n=8 independent, matched healthy human donor rested and stimulated iNKT cells and T_CONV_ (processed per schematic in Supplemental Figure 1) relative expression of fatty acid oxidation (FAO) pathway genes in NanoString nCounter Human Metabolic Pathways probe set. Genes with counts under 100 were eliminated from analysis. Coloring indicates relative expression of each gene, from low (blue) to high (red). Heatmap generated on Morpheus. **(D)** Purified rested and stimulated human iNKT cells and T_CONV_ were stained for intracellular Cpt1a expression after 48 hours. Mean fluorescence intensity (MFI) of Cpt1a (left) and percentage of Cpt1a positive cells (right) for each cell type relative to isotype control indicated. Each symbol represents independent, matched healthy human donor sample. **(E)** Heatmap of n=8 independent, matched healthy human donor rested and stimulated iNKT and T_CONV_ relative expression of AMPK pathway genes in NanoString nCounter Human Metabolic Pathways probe set analyzed as in (C). For all graphs, asterisks indicate statistical significance (* p<0.05, ** p<0.01, ***p<0.001).

In T_CONV_, Cpt1a-mediated long-chain FAO supports the survival of memory T cells and regulatory T cells (Michalek et al., 2011; Pearce et al., 2009). One master regulator that promotes FAO metabolism and memory cell differentiation is the nutrient sensor adenosine monophosphate-activated protein kinase (AMPK), which inhibits mTORC1 to promote catabolism and FAO, particularly in conditions of nutrient stress (Pearce et al., 2009; Rolf et al., 2013). To further investigate whether iNKT cells displayed memory-like metabolism by employing FAO, we interrogated the expression of the 29 annotated genes in the NanoString AMPK signaling pathway probe set in human donor cell subsets. Intriguingly, of the 14 genes significantly differentially expressed between stimulated iNKT cells and T_CONV_, 13 out of 14 were significantly higher in iNKT cells than stimulated T_CONV_ (**Figure 4E**). This further supports an important and distinct role for FAO metabolism in human iNKT cells relative to T_CONV_.

### Stimulated human iNKT cells oxidize fatty acids to a greater extent than stimulated T_CONV_

Given the striking differences in the expression of *Cpt1a* and other FAO genes between stimulated human iNKT cells and T_CONV_, we postulated that these cells may differ in their use of FAO metabolism. To investigate the dependence on FAO for the metabolic activity of iNKT cells and T_CONV_ upon stimulation, we performed Seahorse extracellular flux analysis to generate real-time metabolic measurements of these cells. To specifically determine the contribution of fatty acids to the oxygen consumption rate (OCR) of stimulated iNKT cells and T_CONV_, we injected into the cells either media only (vehicle) or etomoxir, a pharmacological inhibitor of Cpt1a. Upon etomoxir treatment, long-chain fatty acid import into the mitochondria is blocked, such that cytosolic fatty acids cannot be oxidized via the TCA cycle to fuel OXPHOS. Interestingly, we found that upon addition of carbonyl cyanide-4 (trifluoromethoxy) phenylhydrazone (FCCP) – which decouples the mitochondrial membrane to drive maximal substrate demand – etomoxir-treated iNKT cells displayed significantly reduced maximal respiration (**Figure 5A, 5C**). This indicates that fatty acids represent an important substrate for oxidation in stimulated iNKT cells. In contrast, stimulated T_CONV_maintained equal levels of OCR upon FAO inhibition (**Figure 5B-5C**), implying that in these cells, fatty acids do not represent a substrate for oxidation. In addition, we also quantified mitochondrial respiration-linked ATP production, defined as the difference in OCR at baseline and upon injection of oligomycin A, an ATP synthase inhibitor. Notably, this parameter was also significantly lowered in etomoxir-treated iNKT cells, but not in T_CONV_ (**Figure 5D**), further supporting the importance of FAO for contributing to ATP production derived from mitochondrial respiration in stimulated iNKT cells. Interestingly, depletion of fatty acids did not completely ablate OCR activity of iNKT cells, as they still displayed heightened respiration upon addition of FCCP above basal OCR levels. This may suggest that iNKT cells do not solely depend on fatty acids and employ additional metabolic substrates for energy consumption upon stimulation. Nevertheless, our findings ultimately reveal a key bioenergetic difference between stimulated human iNKT cells and T_CONV_, whereby in stimulated iNKT cells, fatty acids serve as a substrate contributing to fueling the TCA cycle and OXPHOS, while T_CONV_ preferentially rely on other substrates for oxidative metabolism, such as glucose and glutamine.

**Figure 5:**
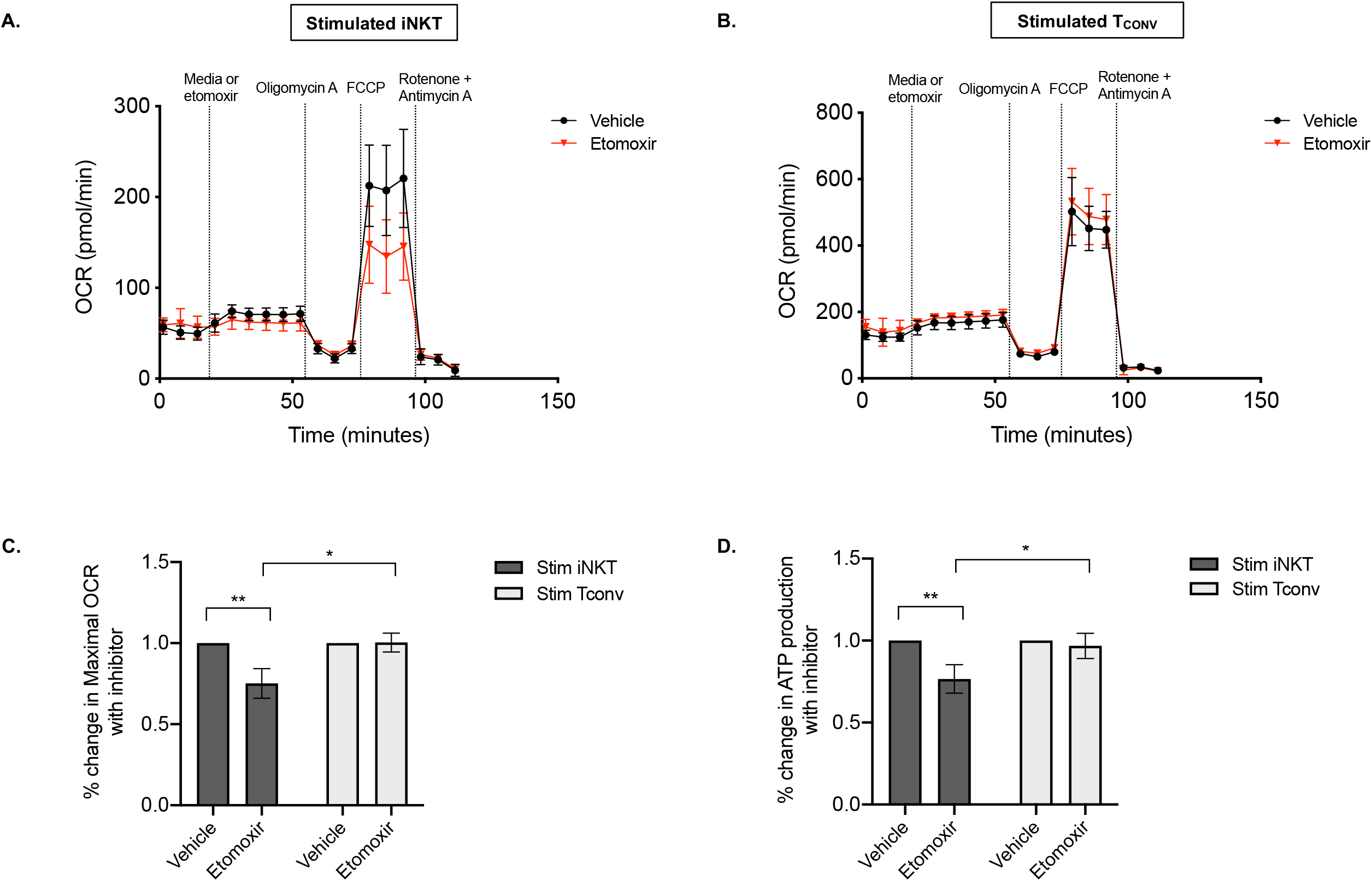
Stimulated human iNKT cells oxidize fatty acids more than stimulated T_CONV_. **(A-B)** Real-time measurements of oxygen consumption rate (OCR) of 48-hour stimulated iNKT cells (A) and stimulated T_CONV_ (B) from matched human donors were obtained on a Seahorse Bioanalyzer. OCR was measured after sequential additions of media (vehicle, black) or 4μM etomoxir (red) followed by 1.5μM oligomycin A, 0.5μM FCCP, and 0.5μM rotenone and antimycin A. Graphs depict representative example of 3 independent, matched human donor replicates. **(C)** Summary data displaying the percent change in maximal OCR upon FCCP (mitochondrial uncoupler) injection with pre-injection of etomoxir relative to vehicle controls for stimulated iNKT cells (dark grey) and stimulated T_CONV_ (light grey). Graphs depict summary data from 3 independent matched human donors. Asterisks indicate statistical significance (* p<0.05, ** p<0.01). **(D)** Summary data displaying the percent change in ATP production with etomoxir addition relative to vehicle controls for stimulated iNKT cells (dark grey) and stimulated T_CONV_ (light grey). Graphs depict summary data from 3 independent matched human donors. Asterisks indicate statistical significance (* p<0.05, ** p<0.01).

Taken together, our data reveals that human iNKT cells possess a unique metabolic profile from T_CONV,_ characterized by greater FAO metabolism and a reduced requirement for glucose and glutamine for anti-tumor effector functions upon activation. Importantly, these differential bioenergetic requirements may allow iNKT cells to retain their functional activity in the nutrient-poor solid TME, where they may possess an advantage in competing with tumor cells relative to T_CONV_ that could potentially be exploited therapeutically.

## DISCUSSION

One of the major limitations of iNKT-cell based immunotherapies is the lack of understanding of the cellular properties enabling long-term persistence within the TME. While new insights have begun to highlight the importance of cellular metabolism for sustained effector function and persistence of effector T cells within the TME, these insights have not been extended to iNKT cells – particularly human iNKT cells. To address this gap in knowledge, we sought to delineate the bioenergetic requirements of human iNKT cells relative to their conventional T cell counterparts. In the present study, we demonstrate for the first time that primary human PBMC-derived iNKT cells possess distinct metabolic features from matched T_CONV_ at both baseline and upon stimulation, and that these could impact anti-tumor effector functions. Specifically, we find that iNKT cells do not depend on glucose and glutamine for anti-tumor cytokine production and cytotoxicity, and rather possess a “memory-like” metabolic phenotype characterized by high mitochondrial mass and fatty acid oxidation metabolism. We believe these novel bioenergetic differences distinguish iNKT cells from T_CONV_ and that these findings could suggest notable differences in the ability of these cells to adapt, survive, and function in nutrient-poor TME conditions – an assertion that has important implications for the use of iNKT cells in cancer immunotherapy.

The TME imposes metabolic challenges to the functions of T cells and other immune cells that directly impact antitumor immunity and tumor progression. As cancer and myeloid cells employ aerobic glycolysis to support biosynthetic requirements for proliferation via the Warburg effect, they rapidly uptake glucose, glutamine, and amino acids, depleting the TME of these nutrients (DeBerardinis and Chandel, 2016; Ghoshdastider et al., 2021; Pavlova and Thompson, 2016; Reinfeld et al., 2021); furthermore, the poor vasculature creates regions of hypoxia within the TME. As such, the hypoxic, acidic, nutrient-deplete TME impairs the ability of T_CONV_ to sustain their functional activity. Indeed, it is well appreciated that the ability of T_CONV_ to engage memory-like FAO metabolism results in improved persistence and anti-tumor activity within the TME (reviewed in Kishton et al., 2017). As such, the design of metabolic interventions to improve the efficacy of solid tumor immunotherapies provide an attractive therapeutic strategy. However, due to the metabolic similarities between tumor cells and effector T_CONV_ (Allison et al., 2017), preserving anti-tumor immune cell function while specifically targeting tumor cell metabolism is difficult. Our observations, however, imply that the distinct bioenergetic requirements employed by iNKT cells may allow for the utility of metabolic modulators to specifically target tumor cells and their supportive myeloid cells without profoundly affecting iNKT cell effector functions. Specifically, we found that stimulated iNKT cells retain anti-tumor functions despite glucose and glutamine depletion; furthermore, unlike T_CONV_, the effector mechanisms of iNKT cells appear to be decoupled from Myc signaling. Since Myc is an oncogenic driver in many cancers, targeting its activity is a key therapeutic strategy for attenuation of tumor cell growth. While such strategies would also significantly inhibit T_CONV_ activation (Wang et al., 2011), iNKT cell activity may be less affected, supporting the notion that adjunctive therapies that both boost iNKT cell function and inhibit tumor metabolism could be therapeutically valuable. Similarly, the use of inhibitors of glucose and glutamine metabolism may effectively target the bioenergetics of tumor cells and tumor-supportive myeloid cells while allowing for sustained iNKT cell anti-tumor functions. Thus, the unique metabolic features of iNKT cells may enable the use of broader metabolic interventions to which T_CONV_ may be particularly sensitive.

Interestingly, we observed that iNKT cells displayed higher expression of AMPK signaling genes relative to T_CONV_. AMPK is a nutrient sensor that inhibits mTORC1 to promote catabolism and mitochondrial metabolism, including FAO (Pearce et al., 2009; Rolf et al., 2013). Importantly, AMPK pathway activity has been shown to promote T cell longevity and survival, antigen recall responses in memory T cells (Blagih et al., 2015; Kishton et al., 2016), and additionally promote T_REG_ differentiation and function (Michalek et al., 2011). Our data suggests that like memory T cells and T_REG_ (as well as additional immunosuppressive populations, such as tumor-associated macrophages and myeloid-derived suppressor cells), the upregulation of AMPK-mediated catabolic metabolic programs may allow iNKT cells to adapt to the nutrient-deplete conditions of the TME. This presents a clear distinction from effector T_CONV_, which depend upon anabolic metabolic pathways to sustain anti-tumor functions. In T_REG_, the Forkhead Box protein (Foxp3) inhibits Myc signaling and glycolysis to promote OXPHOS and allow enhanced function in low-glucose, lactate-rich environments (Angelin et al., 2017). Although lactic acid has previously been shown to blunt the effector functions of T and NK cells (Brand et al., 2016), as well as murine iNKT cells *in vitro* (Xie et al., 2016), a further dissection of its contribution to human iNKT cells and a better understanding of the metabolic flexibility of iNKT cells in acidic TME environments is important. Intriguingly, a recent study demonstrated that intratumoral T_REG_, relative to peripheral T_REG_, require the uptake of lactate – secreted by tumor cells – to maintain their suppressive effector functions within the TME (Watson et al., 2021). It is thus possible that human iNKT cells, given their overlapping metabolic profile with T_REG_, could utilize similar mechanisms by which to employ metabolic flexibility in adapting to the acidic, nutrient-deplete TME. Given the link between *ex vivo* expansion and *in vivo* persistence of T_CONV_ within solid tumors (Kishton et al., 2017), a greater understanding of the properties that would allow long-term persistence of exogenously expanded iNKT cells upon adoptive transfer is desired. Nevertheless, our data suggest that at least some proportion of human PBMC-derived iNKT cells are memory-like in metabolism and may thus prove to be persistent within the TME.

Despite being in its early clinical stages, iNKT cell-based immunotherapies have begun to demonstrate some promise for solid tumors. One approach in phase I and II trials is the direct, adoptive transfer of activated iNKT cells, which has been tested in non-small cell lung cancer (Shin et al., 2001), head and neck squamous cell carcinoma (HNSCC; (Kunii et al., 2009)), and melanoma (Exley et al., 2017); these studies have demonstrated a transient boost in the numbers of circulating iNKT cells in patients and moderate stabilization of disease progression. Another approach currently under clinical investigation is the use of chimeric antigen receptor (CAR)-enabled iNKT cells. CAR-iNKT cells directed towards GD2, a disialoganglioside highly expressed on malignant neuroblastoma cells, are currently in phase I trials and preliminary studies indicate that the adoptively transferred iNKT cells localize to the tumor site and mediate tumor regression (Heczey et al., 2014, 2020).

A remaining challenge for the adoptive transfer strategies using iNKT cells is the ability to effectively expand sufficient numbers of cells *ex vivo*, given the low starting frequencies in peripheral blood. Recently, Zhu et al. demonstrated the preclinical feasibility and efficacy of hematopoietic stem cell (HSC)-derived iNKT cells to induce anti-tumor cytotoxicity in both hematologic and solid tumor models (Zhu et al., 2019). This may allow for greater scalability and broader utility of iNKT-cell based immunotherapies. Additionally, given that iNKT cells are stimulated by monomorphic CD1d, it is formally possible to explore the use of allogeneic off-the-shelf iNKT cell products in hosts with severe immunocompromise who lack endogenous T cells and are incapable of host vs. graft responses. Interestingly, two studies of both *ex vivo*-expanded iNKT cells and CAR-iNKT cells have demonstrated that the most persistent effector populations – that retain anti-tumor function and maintain longevity within the TME – are those that express CD62L (Tian et al., 2016) and are transduced with IL-15 (Xu et al., 2019). Notably, in T_CONV_, IL-15 promotes a more memory-like metabolic profile (van der Windt et al., 2012), and CD62L is also a central memory marker that is correlated with stem-like properties and enhanced anti-tumor efficacy (Graef et al., 2014; Sommermeyer et al., 2016; Wang et al., 2012); together, this further supports the link between iNKT cell metabolism and persistence in a tumor context. Overall, while iNKT cell-based immunotherapy platforms have demonstrated some early promise, ultimately, the long-term persistence and clinical efficacy remains unknown. However, the optimization of these strategies with additional knowledge gained from studies of iNKT cell metabolism would be of great value.

The present study is the first to characterize primary human iNKT cell metabolism side-by-side with T_CONV_, and interestingly, reveals bioenergetic and functional differences between these lymphocyte populations that may bear important future clinical impact. Importantly, while our results suggest that iNKT cells may be more facile at adapting to the TME, they also prompt the need for metabolic characterization of iNKT cell subsets at baseline and upon stimulation directly from within the TME.

## AUTHOR CONTRIBUTIONS

P.K. and H.B. conceived study and wrote manuscript. P.K. designed and performed experiments, analyzed data, and constructed figures. C.B. contributed to conducting experiments and data analysis. S.A.G. and D.M.B. provided key reagents required for experiments. U.H.B. and D.M.B. provided technical expertise, contributed to experimental design and analysis, and critically reviewed manuscript. All authors contributed to the manuscript and approved of the submitted version.

## ACKNOWLEDGEMENTS

We would like to thank our colleagues Sunny Shin, Kathryn Wellen, Taku Kambayashi, and Will Bailis (University of Pennsylvania) for offering scientific expertise in experimental analysis and manuscript preparation. We also thank our former lab member Gabrielle Ferry (University College London) for critical review of the manuscript. We gratefully acknowledge Rajat Das and Ted Hofmann (CHOP) for providing key technical assistance with Seahorse flux metabolic assays and NanoString nCounter transcriptional profiling, respectively. Kevin Bittman (Agilent) provided technical guidance and data analysis support for Seahorse experiments, and Allison Songstad (NanoString) assisted with NanoString data analysis. We would finally like to thank the CHOP Flow Cytometry Core and the University of Pennsylvania (UPenn) Flow Cytometry and Human Immunology Cores for providing key reagents (primary human cells) and instrumentation (cell sorters) required for experiments.

## DATA AVAILABILITY STATEMENT

Data requests may be directed to the corresponding author, Hamid Bassiri, at.

