## Supplementary figures and tables for "Distinct bioenergetic features of human invariant natural killer T (iNKT) cells enable retained functions in nutrient-deprived states"

### Supplementary Material

**Khurana et al. (2021)**

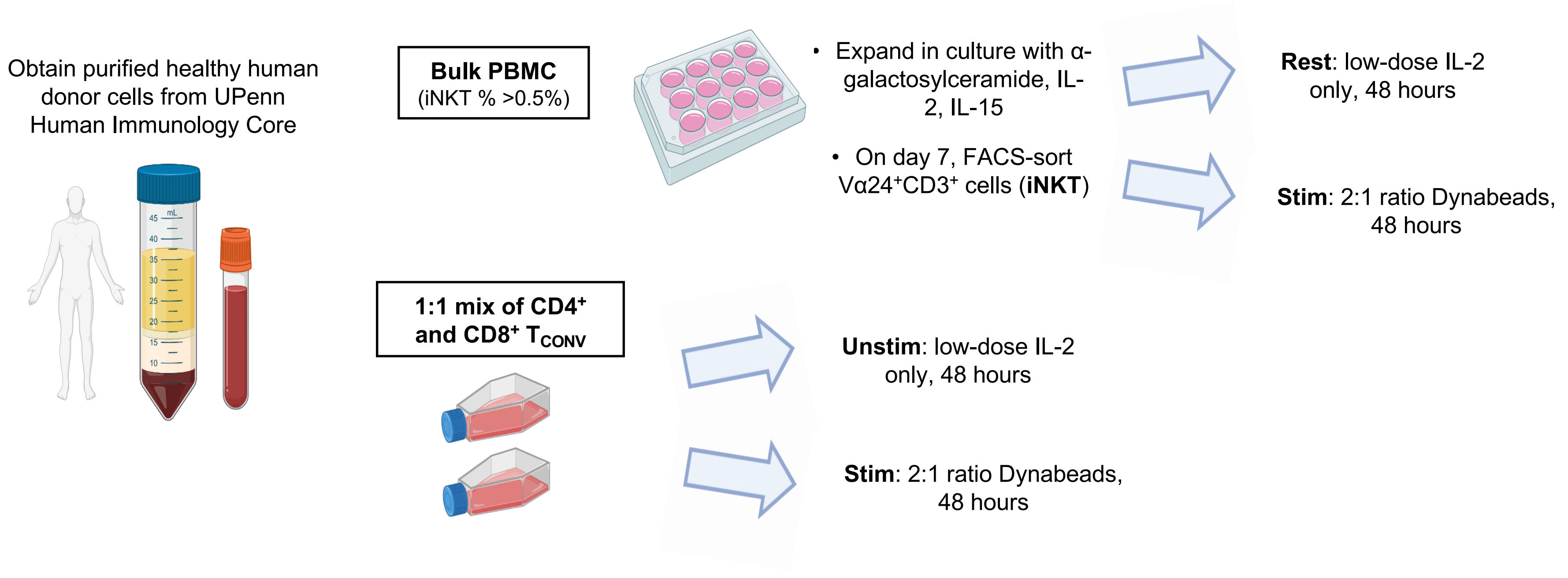


Supplementary Figure 1: **Expansion and stimulation scheme for purifying healthy human donor PBMC-derived iNKT cells and conventional T cells ($T_{CONV}$).** Schematic depicting obtaining and expanding purified human donor cells from UPenn Human Immunology Core. For all studies, iNKT cells were expanded, FACS-sorted, and rested as indicated. $CD4^+$ and $CD8^+$ conventional T cells ($T_{CONV}$) from matched sets of human donors were mixed at a 1:1 ratio to ensure equal composition of these subsets for consistency in all studies. Matched iNKT cells $T_{CONV}$ and were subjected to 48 hours of “rest” and “stimulation” with CD3/CD28 Dynabeads as indicated. Images created using BioRender.

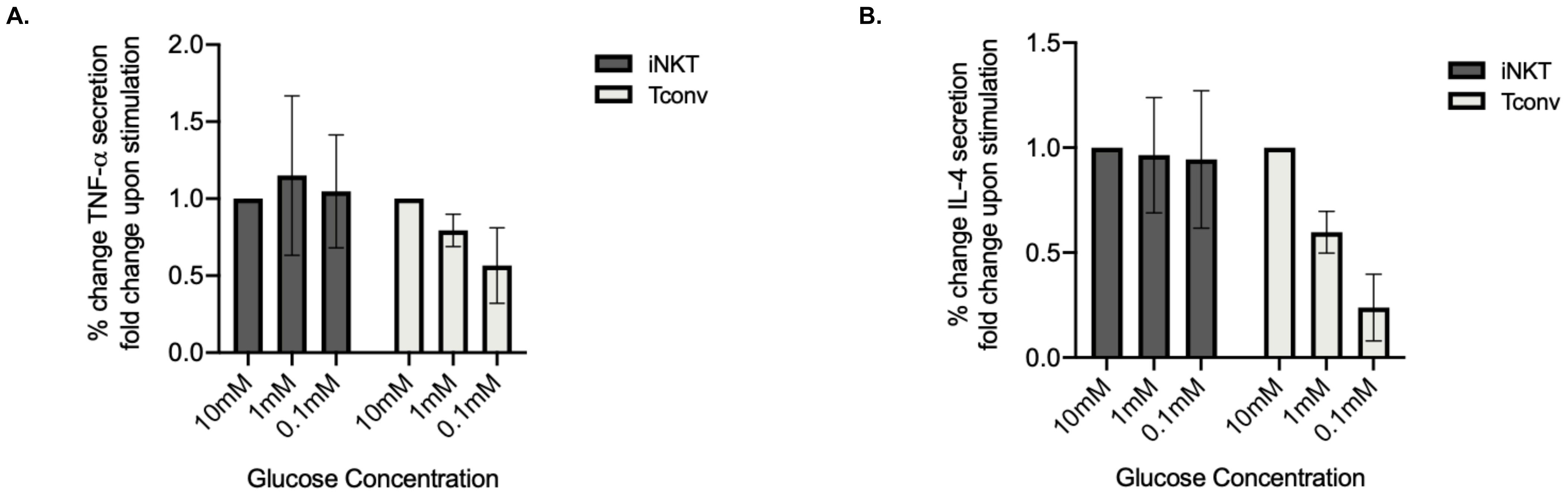


**<u>Supplementary Figure 2</u>: Human iNKT cells maintain production of additional cytokines in glucose-depleted media relative to $T_{CONV}$.** Supernatants were collected from rested and stimulated iNKT cells and $T_{CONV}$ cultured in 10mM, 1mM, or 0.1mM glucose after 48 hours. Secreted TNF-α **(A)** and IL-4 **(B)** were quantified via MSD Multiplex Assay. Summary data of percent change in fold change upon stimulation for 3 independent, matched human donors depicted in bar graphs for stimulated iNKT cells (dark grey) and stimulated $T_{CONV}$ (light grey).

A.

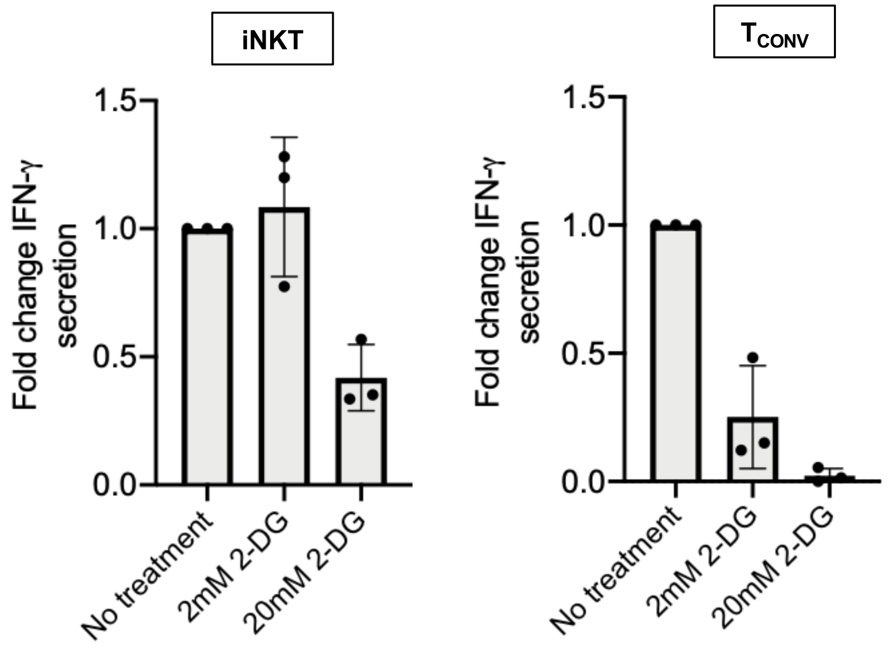


B.

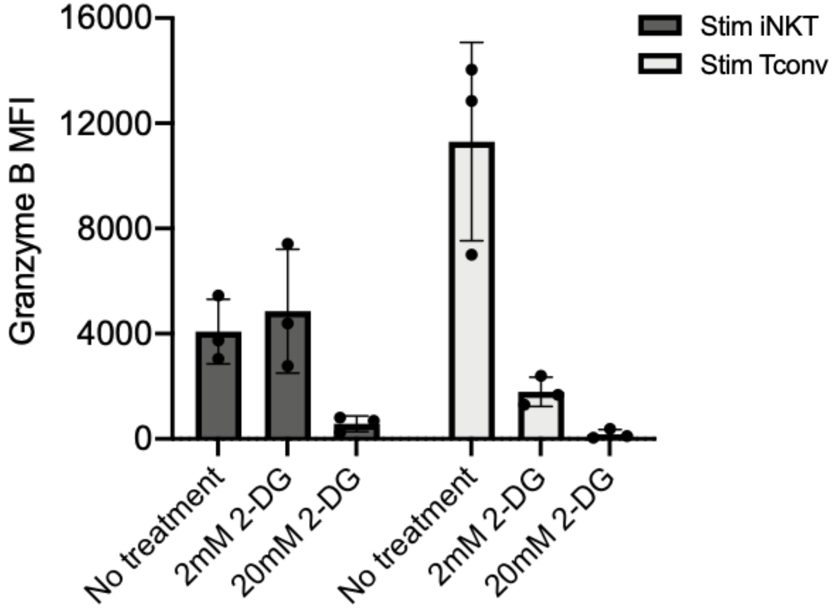


<u>Supplementary Figure 3</u>: **Human iNKT cells are less sensitive to pharmacological inhibition of glycolysis than $T_{CONV}$. (A)** Sorted PBMC-derived iNKT cells and $T_{CONV}$ were rested or stimulated for 48 hours in complete AIM V media with low-dose IL-2 (30U/mL) only or containing either 2mM or 20mM 2-Deoxy-D-glucose (2-DG). Supernatants from rested and stimulated iNKT cells and $T_{CONV}$ cultured in each condition were collected after 48 hours and profiled for levels of IFN-γ via ELISA. Fold change in IFN-γ secretion upon stimulation is displayed for iNKT cells (left) and $T_{CONV}$ (right) relative to rest and unstimulated conditions, respectively. **(B)** Stimulated iNKT cells and $T_{CONV}$ were stained for intracellular Granzyme B or isotype control after 48 hours of indicated treatments. Bar graph depicts mean fluorescence intensity (MFI) of granzyme B relative to isotype control. Each dot represents independent, matched human donor sample.

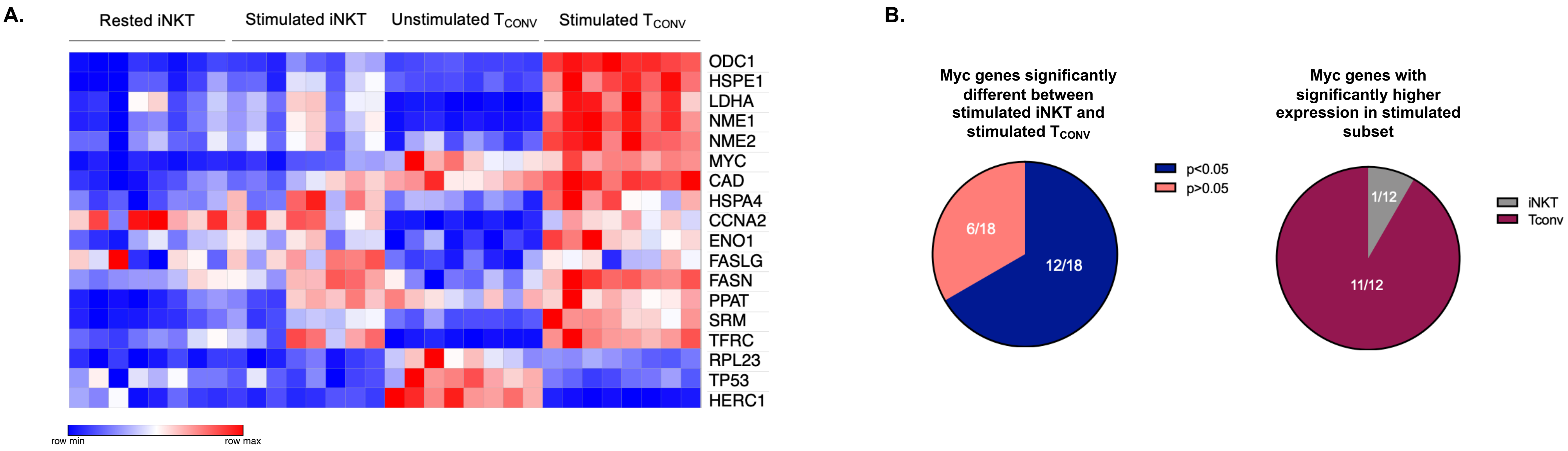


**Supplementary Figure 4: Stimulated human iNKT cells have lower expression of Myc signaling genes than $T_{CONV}$. (A)** Heatmap of n=8 independent, matched healthy human donor-derived rested and stimulated iNKT and $T_{CONV}$ relative expression of Myc pathway genes in NanoString nCounter Human Metabolic Pathways probe set. Genes with counts under 100 were eliminated from analysis. Coloring indicates relative expression of each gene, from low (blue) to high (red). Heatmap generated on Morpheus. **(B)** Pie charts displaying statistical analysis of NanoString Myc pathway genes from (A). Proportions of genes significantly different in stimulated iNKT vs. stimulated $T_{CONV}$ depicted (left), and of those genes, percentages significantly higher in $T_{CONV}$ and iNKT shown (right).

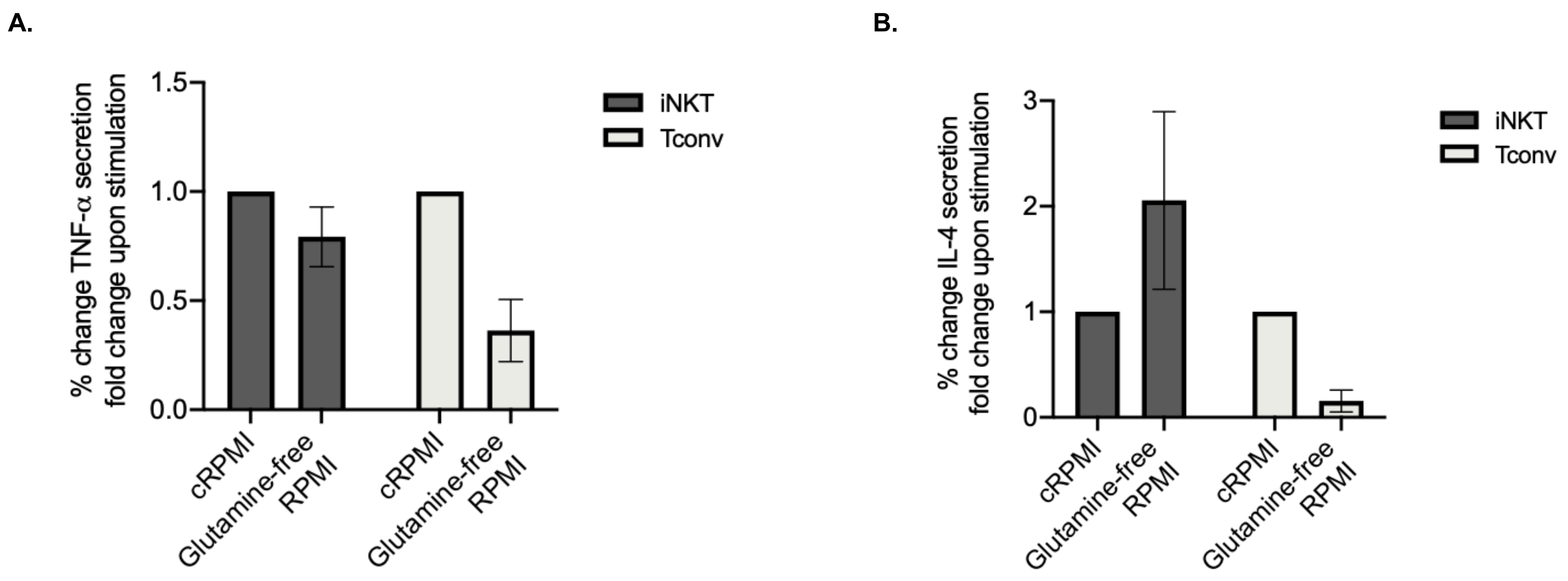


**<u>Supplementary Figure 5</u>: Glutamine is not required for secretion of additional cytokines in human iNKT cells upon stimulation.** Supernatants were collected from rested and stimulated iNKT cells and $T_{CONV}$ cultured in either complete RPMI media or glutamine-depleted RPMI media conditions after 48 hours. Secreted TNF-α **(A)** and IL-4 **(B)** were quantified via MSD Multiplex Assay. Summary data of percent change in fold change upon stimulation for 4 independent, matched human donors depicted in bar graphs for stimulated iNKT cells (dark grey) and stimulated $T_{CONV}$ (light grey).

**Supplementary Table 1. Myc pathway gene set expression in stimulated iNKT vs. $T_{CONV}$**

| Gene name | p-value | q-value | Higher expressing cell subset |
|---|---|---|---|
| ODC1 | 1.7854E-10 | 1.4426E-09 | $T_{CONV}$ |
| MYC | 1.9047E-08 | 7.6950E-08 | $T_{CONV}$ |
| HSPE1 | 1.5100E-06 | 4.0670E-06 | $T_{CONV}$ |
| NME1 | 3.0130E-06 | 6.0862E-06 | $T_{CONV}$ |
| NME2 | 4.3946E-06 | 7.1017E-06 | $T_{CONV}$ |
| CAD | 1.5126E-05 | 2.0369E-05 | $T_{CONV}$ |
| HERC1 | 6.1085E-05 | 7.0510E-05 | iNKT |
| LDHA | 1.2875E-04 | 1.3004E-04 | $T_{CONV}$ |
| SRM | 2.3461E-04 | 2.1063E-04 | $T_{CONV}$ |
| RPL23 | 1.0744E-03 | 8.6815E-04 | $T_{CONV}$ |
| FASN | 1.5111E-02 | 1.1100E-02 | $T_{CONV}$ |
| ENO1 | 1.8666E-02 | 1.2568E-02 | $T_{CONV}$ |
| TFRC | 5.8868E-02 | 3.6589E-02 | $T_{CONV}$ |
| FASLG | 7.2688E-02 | 4.1952E-02 | iNKT |
| PPAT | 1.3568E-01 | 7.3088E-02 | $T_{CONV}$ |
| CCNA2 | 1.8492E-01 | 9.3383E-02 | iNKT |
| HSPA4 | 4.1727E-01 | 1.9833E-01 | $T_{CONV}$ |
| TP53 | 7.4413E-01 | 3.3403E-01 | $T_{CONV}$ |

**Supplementary Table 2. Fatty acid oxidation (FAO) pathway gene set expression in stimulated iNKT vs. $T_{CONV}$**

| Gene name | p-value | q-value | Higher expressing cell subset |
|---|---|---|---|
| CPT1A | 1.8304E-08 | 9.2435E-08 | iNKT |
| ACAA2 | 2.2474E-07 | 5.6746E-07 | iNKT |
| ACAT2 | 1.7960E-05 | 3.0232E-05 | iNKT |
| ACOX1 | 1.1788E-03 | 1.4883E-03 | iNKT |
| ACAT1 | 8.2236E-03 | 8.3058E-03 | iNKT |
| PRKAB2 | 1.9726E-01 | 1.6603E-01 | iNKT |
| HADH | 2.3673E-01 | 1.6985E-01 | iNKT |
| PRKAG2 | 2.6907E-01 | 1.6985E-01 | $T_{CONV}$ |
| ECHS1 | 9.4583E-01 | 5.3072E-01 | iNKT |
